# Ex-vivo expansion of patient-derived PBMCs preferentially results in effector memory T-cell proliferation with restored autologous efficiency in HBV-HCC

**DOI:** 10.1101/2024.12.01.626262

**Authors:** Janine Kah, Lisa Staffeldt, Gregor Mattert, Tassilo Volz, Maximilian Voß, Kornelius Schulze, Asmus Heumann, Maura Dandri, Stefan Lüth

## Abstract

**Background:** T-cell-based therapies achieved milestones in targeting solid tumors, such as hepatocellular carcinoma, by leveraging the cytotoxic potential of effector memory T-cells. However, a key challenge lies in the ex-vivo expansion of functional memory T-cells while simultaneously preventing over-differentiation into senescent TEMRA cells.

**Methods:** PBMC derived from 10 HCC patients and two healthy donors were used for expansion in small bioreactors equipped with a permeable membrane for 2 weeks. During expansion, surface marker composition, cytokine and chemokine production were observed. Enriched T-cell subsets from three patients with chronic HBV infection were analysed in detail. We extracted the enriched complement of one chronically infected patient to elucidate the therapeutic potential.

**Results:** We successfully expanded the effector T-cell subsets for all investigated samples and consistently enriched the cell amount over time. Subsequently, we showed that the expanded patient-derived T-cells showed functionality against autologous liver cancer-derived cells by inducing receptor and protein-mediated cell death.

**Conclusion:** We showed the consistent ex-vivo expansion of T-cell subsets of initial patient-derived PBMCs. The enriched subsets exhibit cytotoxic functionality and shift to cytolytic TEMRA cells in an immune evasion setting in the context of chronic HBV-infected patient-derived liver cancer cells.

## Introduction

The progress of T-cell-based therapies reached significant achievements in treating solid tumors by offering the unique advantage of directly targeting cancer cells [1, 2]. Multiple studies have evaluated effector memory T-cells (TEM) and naïve T-cells (TN) as essential elements for successfully eliminating cancer cells [3, 4]. TN cells demonstrate long-lasting persistence and differentiation potential, providing a continuous source of effector T-cells [4, 5]. In contrast, TEM cells respond rapidly to antigens and can retain long-term memory [6, 7]. Terminally developed effector memory T-cells re-expressing CD45RA (TEMRA) are a vital component of the therapeutic T-cell repertoire, notwithstanding TEM cells. These highly cytotoxic yet less proliferative cells are prone to senescence, restricting their application in therapies necessitating long-term persistence [8].

Achieving sufficienT-cell quantities while maintaining optimal differentiation status of the therapeutic complement is a primary problem in T-cell-based methodologies [9–11]. This work aims to preserve the functional integrity of the effective T-cell complement during the enrichment of patient-derived PBMCs taken from persons with various forms of hepatocellular carcinoma (HCC). The immunosuppressive and extremely heterogeneous tumor microenvironment (TME) of hepatocellular carcinomas (HCCs) presents a substantial obstacle for T-cell treatments, rendering the proliferation of effective and functional tissue-resident memory T (TEM) and naïve T (TN) cells a crucial element of tailored immunotherapy [12]. Consequently, our principal objective was to illustrate that we can reliably expand TEM cells from the depleted T-cell subsets of HCC patients. The second objective was to investigate the efficacy of the augmented therapeutic complement in an autologous context against liver cancer cells derived from the same patient. To emphasise the effectiveness of enriched T-cells, we utilised a highly suppressive tumor microenvironment by employing patient-derived LC14 cells obtained from an HBV-related hepatocellular carcinoma [13, 14].

## Methods

### Sample collection and purification

Human blood samples were collected, isolated, preserved and ethically approved as described previously [15].

### Grex scaling

Thawed PBMCs were cultivated on a G-Rex24 Well Plate [16] (Wilson Wolf Manufacturing, Minnesota, USA) using 8 ml CTS™ OpTmizer™ (Gibco, Thermo Fisher, Massachusetts, USA) with 10% human AB Serum, 1% Sodium Pyruvate (100 mM) (Gibco, Thermo Fisher, Massachusetts, USA) and 1% L-Glutamin (200 mM) (Gibco, Thermo Fisher, Massachusetts, USA) per Well. 400 IU/ml rh IL-2 (Sartorius, Göttingen, Germany) and 25 µl CD3/CD28 Dynabeads™ (Gibco, Thermo Fisher, Massachusetts, USA) were added for T-cell activation. The PBMCs were seeded in five differenT-cell counts (3x105, 5 x105, 1 x106,2 x106 and 3 x106 cells). On day 3, 7, 10 and 14, 6 ml supernatant was removed, the cells were counted and harvested for flow cytometry analyses and fresh medium with IL-2 was added.

### Flow cytometry

Flow cytometry analyses of PBMC fractions and enriched T-cells during G-Rex scaling were performed on Celesta™ (panel S9) and SymphonyA3™ (panel S10) (Becton Dickinson, New Jersey, USA). The cells were labeled using the antibodies as listed in **Supplementary Table 1**, (all from Becton Dickinson, New Jersey, USA). The antibodies were incubated for 30 minutes at room temperature, and protected from light. Prior to the staining, the cells were incubated with Human BD Fc BlockTM (Becton Dickinson, New Jersey, USA). The gates were defined using fluorescence minus one (FMO) control.

### Isolation of Oligonucleotides

RNA was extracted from enriched T-cells, LC14 cell lines and spheroids using the RNeasy Mini™ and Micro™ RNA purification kit (Qiagen) [17].

### Measurement of gene expression level using TaqMan-based PCR

For measurement of gene-expression, 2-step PCR was performed. Therefore, complementary DNA (cDNA) synthesis was conducted by using MMLV Reverse Transcriptase™ 1st-Strand cDNA Synthesis Kit (Lucigen, Middleton, Wisconsin, USA) to synthesise RNA complementary DNA, according to the manufacturer’s instructions. Human-specific primers from the TaqMan Gene Expression Assay System, listed in **Supplementary Table 2**, were used to determine gene expression levels (Life Technologies, Carlsbad, California, USA). Samples were analysed using the Quant Studio 7™ Real-Time PCR System (Life Technologies, Carlsbad, California, USA). The human housekeeping gene ribosomal protein L0 (RPL0) was used to normalise human gene expression levels.

### Virological load measurement

HBV DNA was extracted from serum samples with the QiAmp MinElute™ Virus Spin kit (Qiagen, Hilden, Germany). For quantification, TaqMan PCR was performed using an HBV-specific probe, as listed in Supplementary Table 2, and cloned HBV-DNA references were amplified in parallel to establish a standard curve for quantification.

### Cell culture and spheroid formatio

Patient-derived cells were generated as described previously [15]. For 3D conditions, different amounts of LC14 cells were seeded on BIOFLOAT™ cell culture plates in DMEM containing L-glutamine and glucose, supplemented with 1% P/S, 10% Gibco fetal bovine serum (FBS; all from Thermo Fisher Scientific, Waltham, USA) and were obtained for 3 weeks to follow the spheroid formation. Spheroids were used for cu-culture experiments after 3 weeks of formation. LC14 cells stably transduced with the vector LeGO-iG2-Puro+-Luc2 (3rd generation HIV1-derived self-inactivating vector) were used for visualisation. Stable transduction was performed as described previously [15]. During treatment, supernatant and/or T-cells were collected for analysis, and captures were obtained manually or automatically using the BX-780 Microscope (Keyence, Osaka, Japan). For 2D conditions, LC14 were seeded as described previously [15] and used for RNA isolation after forming a monolayer of over 90% density.

### IFN-γ quantification

ELISA targeting IFN-γ were conducted using the ELISA MAX™ Standard Set Human IFN-γ (Biolegend, California, USA). The procedure was done according to the manufacturers protocol with slight alterations. Streptavidin-PolyHRP20 (SDT GmbH, Baesweiler, Germany) was used instead of the peroxidase contained in the Kits. The Streptavidin-PolyHRP20 was incubated for 30 minutes at room temperature on a shaker.

### Cytokine and Chemokine Quantification

The LEGENDplex™ Human Inflammation Panel 1 (13-plex) (BioLegend, San Diego, CA) was used to quantify 13 pro-inflammatory cytokines and chemokines (IL-1β, IFN-α2, IFN-γ, TNF-α, MCP-1, IL-6, IL-8, IL-10, IL-12p70, IL-17A, IL-18, IL-23, IL-33) in cell culture supernatants. Samples (n = 2 technical replicates) were incubated with antibody-conjugated beads, followed by detection with biotinylated antibodies and a Streptavidin-PE label. Data acquisition was performed on the SymphonyA3™ system (Becton Dickinson, New Jersey, USA), and concentrations were calculated using standard dilutions in the LEGENDplex™ Data Analysis Software. For relative quantification, supernatants from untreated controls or cell culture mediums were used.

### Statistics and Sample size

For graph design and statistical analysis, the analysis software GraphPad Prism Version 9 (GraphPad Software, Inc., La Jolla, CA, USA) was used. For 2D RNA isolation, n = 3 wells were used and pooled for the lysis. For cDNA analysis using TaqMan assays, n = 3 technical replicates were used. Expression comparison was analysed using a two-tailed unpaired t-test of the 3 technical replicates for ACT treated against untreated controls (**Figure 5H**). For IFN-γ analysis, supernatant from the indicated samples (n = 3) was collected and pooled for cytokine determination using 2 technical replicates. For Cytometric analysis, samples from biological replicates were pooled and divided into n = 2 replicates for each staining panel (**S10, S9 and Legendplex multiplex analysis**). Panels were measured and analysed separately. Statistical outputs are indicated in the figure legends; p-values were plotted in the graph as follows: * p < 0.05; ** p ≤ 0.01 and *** p ≤ 0.001, **** p ≤ 0.0001.

## Results

### Patient-derived T-cells effectively increase CD8+ T-cells in proportion to CD4+ T-cells and produce elevated levels of inflammatory cytokines

As illustrated in **Figure 1A**, patients with respectable hepatocellular carcinomas were included in the study. Blood withdrawal and isolation were conducted on the day of the primary tumor resection. The patient-derived blood cells were stored in liquid nitrogen for one year and utilised for this investigation following thawing and incubation of the PBMC for three days in a standard six-well plate with the specified medium. On day three, PBMCs were enumerated and subsequently put into the tiny bioreactors for observation, as depicted in **Figure 1**. In B, the fold change in cell numbers at the observation time points and the absolute cell counts (**C**) of patient- and healthy donor-derived PBMCs exhibit an initial expansion rate ranging from 4-fold to 52-fold rise within the first four days in the bioreactor. The previously elevated proliferation rates seemed to diminish in the subsequent days.

**Figure 1:**
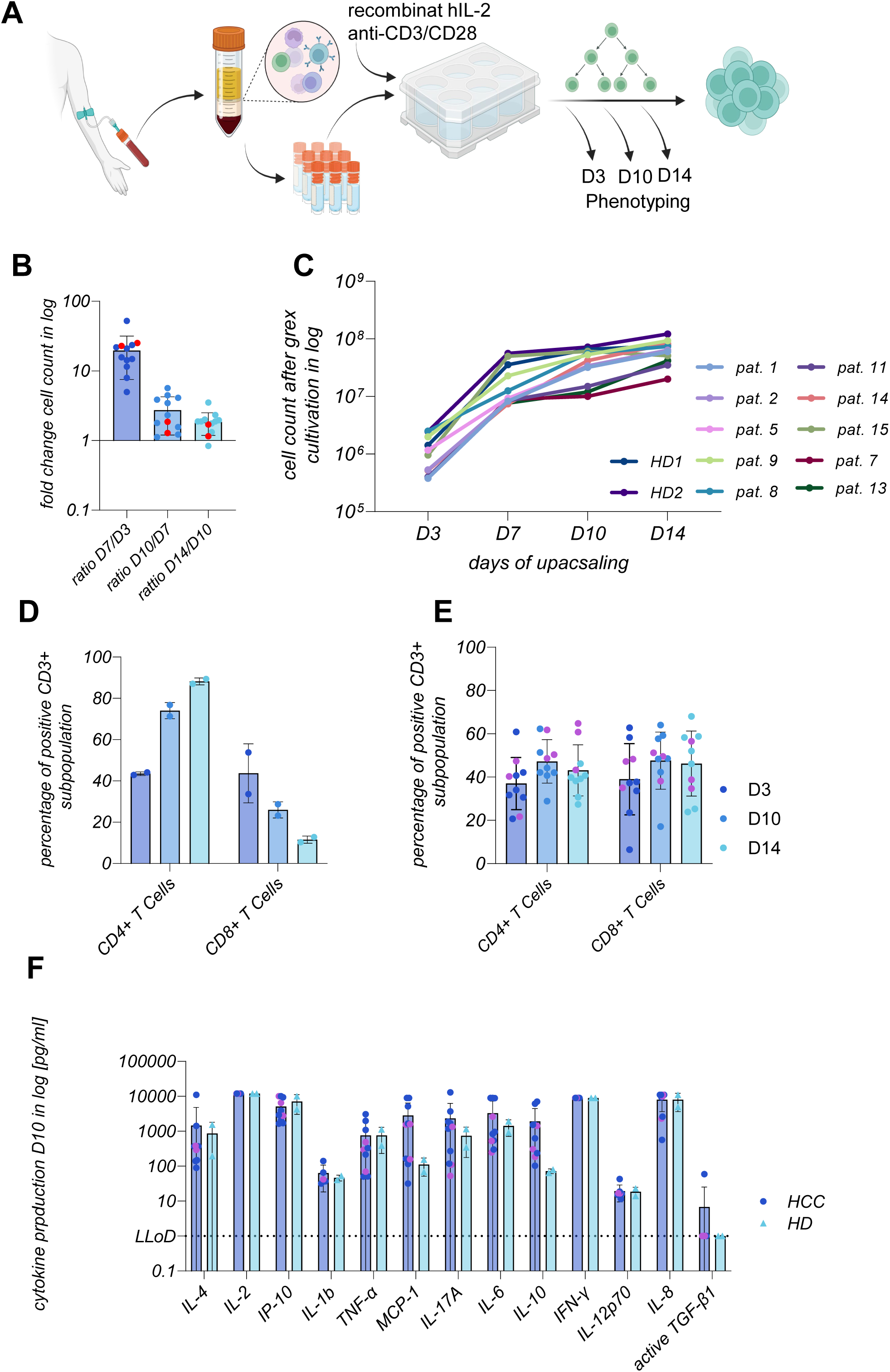
Characterization and classification of patient-derived cells from HBV-related HCC. Figure 1A illustrates the study design. Blood withdrawal from patients with developed HCCs was performed on the day of resection. PBMCs were isolated and cryopreserved before expansion experiments in small G-Rex® bioreactor plates were performed. At indicated time points, immune cells were used for phenotyping. The fold change of the cell counts within the longitudinal growth is illustrated in Figure 1B. Healthy individuals were marked with red dots. Figure 1C shows the longitudinal follow-up of the expansion process for days 3, day 7, day 10, and 14. For the expansion process, PBMCs derived from HCC patients with different HCC-related entities, ages (30-90y) and gender (f/m) were used in direct comparison to PBMCs isolated with the same protocol from healthy donors (HD#1, HD#2). In Figures 1D **and E**, flow cytometric analysis of CD4 and CD8 positive CD3 presenting T-cells is shown during the scaling process. In Figure 1D, healthy donor-derived PBMCs were presented; in Figure 1E, we extracted the CD4/CD8 ratio for all observed patient-derived samples. In Figure 1F, the cytokine and chemokine signature was presented during the expansion, whereby the production of cytokines and chemokines was compared between the healthy and patient groups. Purple dots represent patients with HBV-related HCC. Single dots in the graphs represent individuals. All fractions were presented in percentage of parents and were displayed using mean - +SD.

Nonetheless, all patienT-cells proliferated to substantial cell quantities. To trace the distribution of cell-specific markers and thus identify the cell types in the proliferating culture, we obtained samples for characterisation, as illustrated in **Figure 1D and E**. Cells obtained from HCC patients demonstrate a distinct equilibrium between CD4 and CD8 expressing T-cells throughout time, in contrast to those derived from healthy individuals. The disproportionate growth of healthy individuals results in a CD4:CD8 ratio of 2.5:1 after 14 days of expansion. **Figure 1E** illustrates that patient-derived CD3 T-cells were consistently observed in a 1:1 ratio between CD4 and CD8, with particular instances showing a preferential enrichment of CD8 positive subsets over time. **Figure 1F** illustrates the cytokine and chemokine profile of the expansion culture assessed on day 10. Most analytes were produced in comparable quantities from healthy and patient-derived T-cell subsets, highlighting the pro-inflammatory and migratory characteristics of the expanded populations. The elevated production of the immunosuppressive cytokine IL-10 highlights the source of the patient’s immune cells, indicating a partially immunosuppressive characteristic of the T-cell population.

### The enhancement of T-cell complements from various donors results in elevated populations of TEM with diminished inhibitory signals

Expanded T-cell subsets obtained from HCC patients were evaluated for T-cell differentiation levels and the expression of inhibitory markers. **Figure 2A** illustrates the differentiation of the CD45/CD3 cell subset into TN-like, TCM, TEM, and TEMRA cells based on the expression of CD45RA and CCR7. TN-like cells were analysed for CD95 expression to distinguish TN cells. **In Figures 2B and D**, PBMCs from healthy individuals were differentiated into the specified cell subtypes, whereas in **Figures 2C and E**, PBMC specimens from HCC patients were represented for the same subsets. HD- and HCC-derived PBMCs increased the proportion of TEM cells in the CD4 and CD8 subsets of CD3 T-cells. Cells taken from both healthy donors and HCC patients demonstrate a consistent pattern in the proliferation of CD4 T-cells, characterised by a reduction in the proportions of TN, TCM, and TEMRA cells, next to an increase in memory effector T-cells across the observation period. Upon examining the CD8 positive fraction, we observed that HD displays a similar pattern to that identified in CD4 T-cells, as illustrated in **Figure 2D**. **Figure 2E** illustrates that the patient-derived CD8 T-cells exhibited elevated proportions of TEMRA cells on day 3 post-initiation. The differentiation status of highly cytotoxic CD8 T-cells was nearly absent by day 14. The presence of inhibitory proteins was additionally identified and represented as fold change relative to the baseline samples, illustrated in **Figure 2F** (CD4 TEM) and **Figure 2G** (CD8 TEM). In both fractions, most patient-derived TEM exhibit unchanged or diminished regulatory surface protein expression. In the healthy individuals (red dots), we observed no induction in expression, except for the CD279 expression on CD8 TEM cells in HD#1, as illustrated in **Figure 2G**. In the patient cohort, CD152, CD272 and TIGIT expression elevated heterogeneously, as shown in **Figures 2F and G**. In patient#15, who had the lowest growth rate on day 14, as illustrated in **Figure 1B**, we observed the highest induction of TIGIT and CD152 in both fractions. We observed a reduced expression of the specified inhibitory proteins in 60% of the CD4 and 50% CD8, an unchanged expression in 17% of both and an increase in 23% of the CD4 and 33% of the CD8 TEM cells, whereby 17% of the CD4 TEMs and 7% of the CD8 TEMs reached a 2-fold or higher increase of the inhibition.

**Figure 2:**
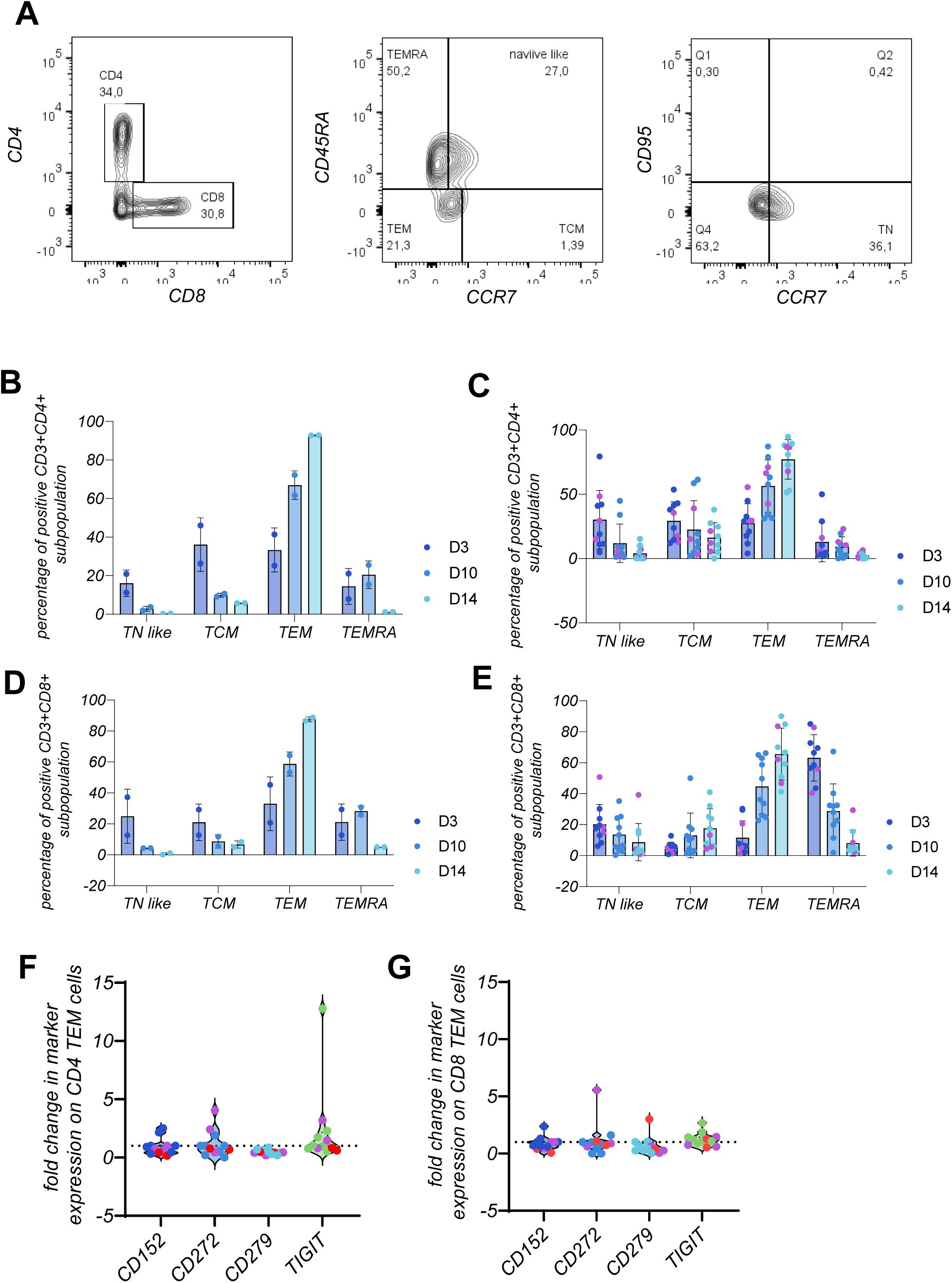
Longitudinal distribution of T-cell subsets in CD4 and CD8 fractions during expansion in patient-derived PBMC. Figure 2A shows the detection of T-cell subsets using flow cytometry. As shown, CD4 and CD8 T-cell subsets were used for the detection of CD45RA and CCR7 to differentiate into a) CD45RA+ CCR7+ (TN); b) CD45RA- / CCR7+ (TCM); c) CD45RA- / CCR7- (TEM) and d) CD45RA+ / CCR7- (TEMRA) cells, as mentioned in the panel in Figure 2A. Figure 2B – E summarizes the extracted T-cell phenotypes over time and distinguishes them into CD4 (B, C) and CD8 (D, E) T-cell fractions from patients (C, E) and healthy donors (B, D). Figures 2F **and G** extracted the TEM fraction on day 14, which was most prominent in all observed up-scaled specimens, to visualize the inhibitory signature for CD152, CD272, CD279 and TIGIT in fold change to the initial status on day 3. In Figure 2F, the TEM CD4 fraction is displayed, and in Figure 2G, the TEM CD8 fraction is indicated. For the comparison analysis, we used patient PBMC n = 10 and healthy donor PBMC n = 2 to calculate mean +-SD, as shown by the error bars in **panels B – E**. In **panels F and G**, we used floating bars to visualize the detected range and the mean of the marker expression.

Despite the finding that all T-cell subsets exhibited high TEM levels, the identified immune signature underscores the significant heterogeneity of the underlying disease, thereby emphasising the necessity of comprehending the individual T-cell composition for effective cellular immunotherapy. Considering this, we examined the particular patterns of three patients clinically characterised as HBV-infected HCC patients eligible for resection. We assessed this characterisation by detecting HBV DNA, as illustrated in **Figure 3A**.

**Figure 3:**
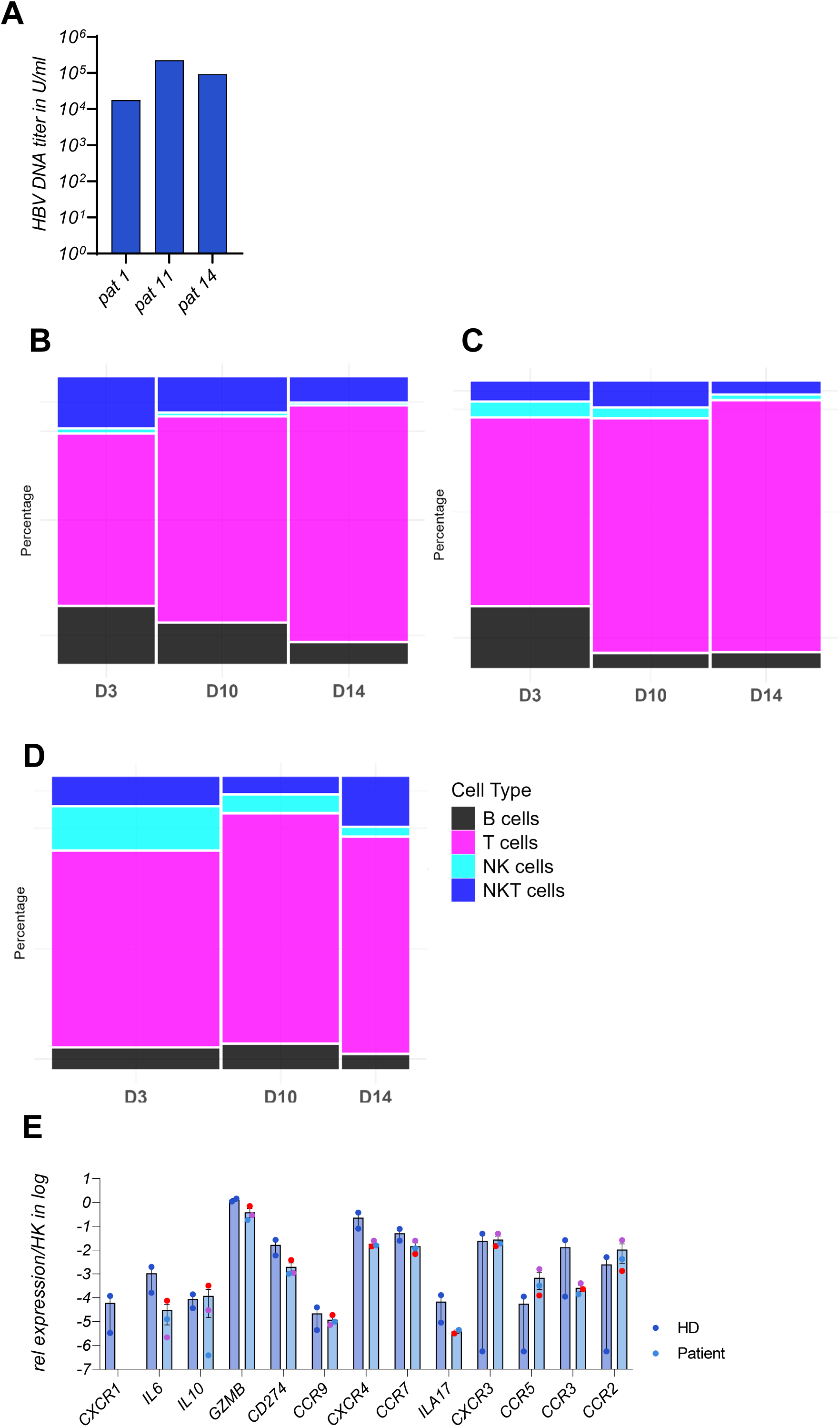
Distribution of immune cell population from patients with HBV-related HCC over time. Figure 3A shows the HBV DNA titer in the plasma derived from the 3 patients on the day of resection, plotted in IU/ml. **Figures 3B – D** show the immune phenotyping for 3 patients with HBV-related HCC over time (B = pat#1, C = pat#11, D = pat#14). As indicated in Figure 3D, main immune cell types were plotted in the percentage of CD45+ / vital cells over time during expansion. Datasets were plotted in a mosaic diagram, whereby each bar of each timepoint represents the part of the whole (=100%), and the diameter means the total amount of CD45+ / vital cell per time point and patient compared to the other. In Figure 3E, up-scaled cell fractions were used for gene expression analysis after 14 days of proliferation and cell differentiation. The results are shown in Figure 3E, normalized against the housekeeper RPL0 and GapDH. Red dots represent pat#1, blue dots represent pat#11, and purple dots represent pat#14. Expression is shown on a logarithmic scale and compared to the gene expression detected in healthy donor T-cells after the same proliferation time. Genes associated with T-cell functionality were observed in mean +- SD.

### Peripheral immune cells in HBV-infected HCC patients exhibit a distinct growth profile

T-cell-based immunotherapy for solid tumors is a personalised approach that must consider the patient’s immunological signature to prevent treatment failure or the onset of cytokine release syndrome during therapy. Consequently, the advancement of personalised medicines will become the benchmark. This section of the study examines the distribution of cell types throughout the expansion phase in the bioreactor for three HBV-positive patients from our cohort. The primary observation was that the distribution of immune cells was highly heterogeneous despite CD3 T-cells being the most proliferative fraction across all patients, as illustrated in **Figures 3B, C, and D**. Notwithstanding this discovery, the presence of NK and NK-T-cells was notable and consistent in patient #14 (D). In patient #1 (B), we observed a significant proportion of NK-T-cells and B cells, which progressively diminishes over time due to the proliferation of CD3 T-cells. In patient #11, a distinct population of NK cells, NK-T-cells, and B cells was observed on day 3, which was also identifiable on day 14 in an equivalent distribution. For patient #1 and patient #11, we observed that the distribution of cell types was analogous to the beginning condition. Notably, for patient #14, NK-T-cells were more pronounced in the day 14 fraction compared to the initial day 3 fraction. In conclusion, the expanded fractions from the studied patients indicate that the T-cell fraction preferentially enriches while maintaining the composition of the starting cell types.

Subsequently, we examined the gene expression levels of the three patients in comparison to the healthy controls on the final day of expansion, day 14, as illustrated in Figure 3E for the specified genes. We observed a complete reduction of CXCR1 gene expression in the three patients relative to the controls. We observed reduced expression of IL-6, GZMB, CD274, CXCR4, and ILA-17 in the three patients, but CCR5 and CCR2 exhibited modestly elevated expression levels. The CCR7, IL-10, CXCR3, and CCR9 expressions were analogous to those in healthy control cells. The expression profile of healthy controls exhibits more significant variability for the cytokine receptors CCR2, CCR3, CCR5, and CXCR3, indicating a particular modification of the complement during the expansion phase. The pro-inflammatory and regulatory profiles were analogous after 14 days of growth, with a marginally elevated migratory and regulatory signature observed in the healthy controls.

In **Figure 4**, we meticulously assessed the immunological condition of the three enlarged complements by extracting patient-specific cytokine and chemokine quantification measurements presented in Figure 1E and T-cell differentiation subsets illustrated in Figures 2C and 2E. The cytokine and chemokine levels of the three patients exhibited a nearly identical profile between patient #1 and patient #14 PBMC. However, PBMCs derived from patient #11 demonstrate a more pro-inflammatory condition. The figures below illustrate the T-cell differentiation status of three patients individually. Patient #11, depicted in **panel C**, exhibits many CD8 TEMRA cells on day 14. This discovery also corresponds with the results of the cytokine release experiment. Notably, patient #14 shows a significant prevalence of TN-like CD8 cells, a characteristic absent in the other patients throughout the expansion.

**Figure 4:**
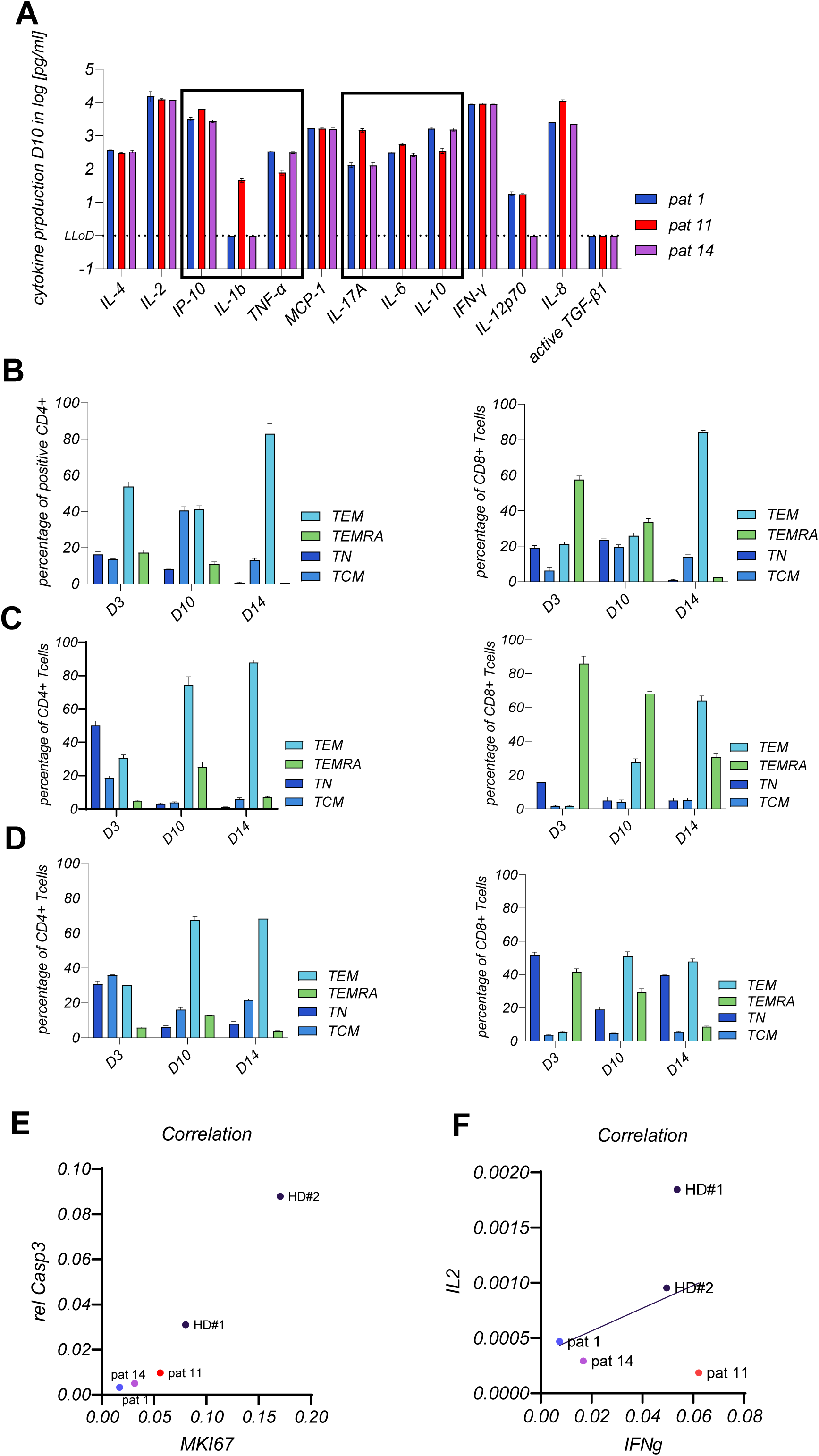
Progression of T-cell differentiation during proliferation of PBMCs derived from HBV-induced HCC patients. Figure 4A plots the patient-specific signature of chemokines and cytokines on a logarithmic scale. Figures 4B-D show the patient-specific follow-up during expansion (B = pat#1, C = pat#11, D = pat#14). The T-cell differentiation subtypes are shown in percentages of the CD4-T-cells (left panel) and the CD8-T-cells (right panel). As mentioned, the T-cells were characterized as TN, TCM, TEM and TEMRA cells. Gene expression levels of proliferation marker ki67 and apoptosis marker caspase 3 were correlated, comparing healthy donors and T-cells derived from HBV-HCC patients. In Figure 4F, the correlation between IL2 and IFN-γ expression is shown for the identical specimens as shown in E. Pearson correlation is plotted on a linear scale.

Furthermore, the gene expression profile depicted in Figure 3E indicated that patient #14 had the highest levels of all examined cytokine receptors. Subsequently, we examined the cellular turnover of the patient’s immune cell complement compared to healthy controls on day 14, utilising gene expression analysis of CAS 3 and MKI67, as illustrated in **Figure 4E**. We detected a positive connection between the expression of MKI67 and CAS3, indicating that the proliferation of immune cells has reached a plateau phase. The healthy controls, particularly HD#2, have a significant turnover of immune cells. The three individuals, conversely, exhibited elevated levels of MKI67, particularly patient #11. This study indicated that the immune cells of the examined patients strive to adapt to the IL-2 environment, resulting in a longer lifespan in the expansion chamber compared to the PBMC of healthy donors. By analysing the expression profiles of IL-2 and IFN-γ to assess the functionality of the enlarged fractions, patient #14 exhibits the optimal balance in the expression of IL-2 and IFN-γ.

### A patient-specific expanded immune cell complement is capable of exerting cytolytic functionality against the patients-derived HCC cells

Considering the cellular turnover, the generation of pro-inflammatory and immunosuppressive cytokines and chemokines, and the identified T-cell subsets, we selected the expanded patient #14 PBMC-derived T-cells to conduct a functionality test with the primary cancer cells generated from patient #14. Consequently, we performed co-culture studies, as illustrated in **Figure 5A**. Patient-derived LC14 cells were cultured under two-dimensional and three-dimensional conditions. They were used for T-cell transfer for 4 and 8 days, respectively, with an expanded immune cell complement at a 1:1 effector-target ratio. **Figure 5B** illustrates a decrease in the CD4 T-cell fraction alongside a notable increase in the CD8 percentage of CD3 T-cells following four days of co-culture. The CD8 fraction exhibited a noteworthy rise in cytolytic TEMRA cells, but the proportions of TEMs and naïve-like cells diminished, as illustrated in **Figure 5C**, left panel. In the CD4 fraction, we observed a significant decrease in TEM cells alongside an increase in the cytolytic TEMRA cell fraction, similar to the CD8 fraction. The transition to TEMRA cells is likely achieved through the continued differentiation of TEMs. This proposal would emphasise the decrease of CD4 T-cells resulting from the brief lifespan of TEMRA cells. The elevated proportion of naïve-like CD8 T-cells in the expanded complement facilitates their differentiation into efficient and cytolytic TEM and TEMRA CD8 cells, increasing CD8 T-cell numbers due to the proliferation of the TEM differentiation status. Analysis of the inhibitory molecules in the subsets depicted in **Figure 5D** (CD4) and E (CD8) revealed a significant increase in the complement, indicated by fold change relative to the baseline, in CD152 across all fractions and subsets after 4 days, particularly in CD4 cells. TIGIT was elevated in all subsets and fractions, except TEMRA cells, while CD279 was predominantly downregulated. CD8 TEM cells exhibited elevated expression of TIGIT, CD152, and CD272 following four days of exposure to patient-derived primary cells.

**Figure 5:**
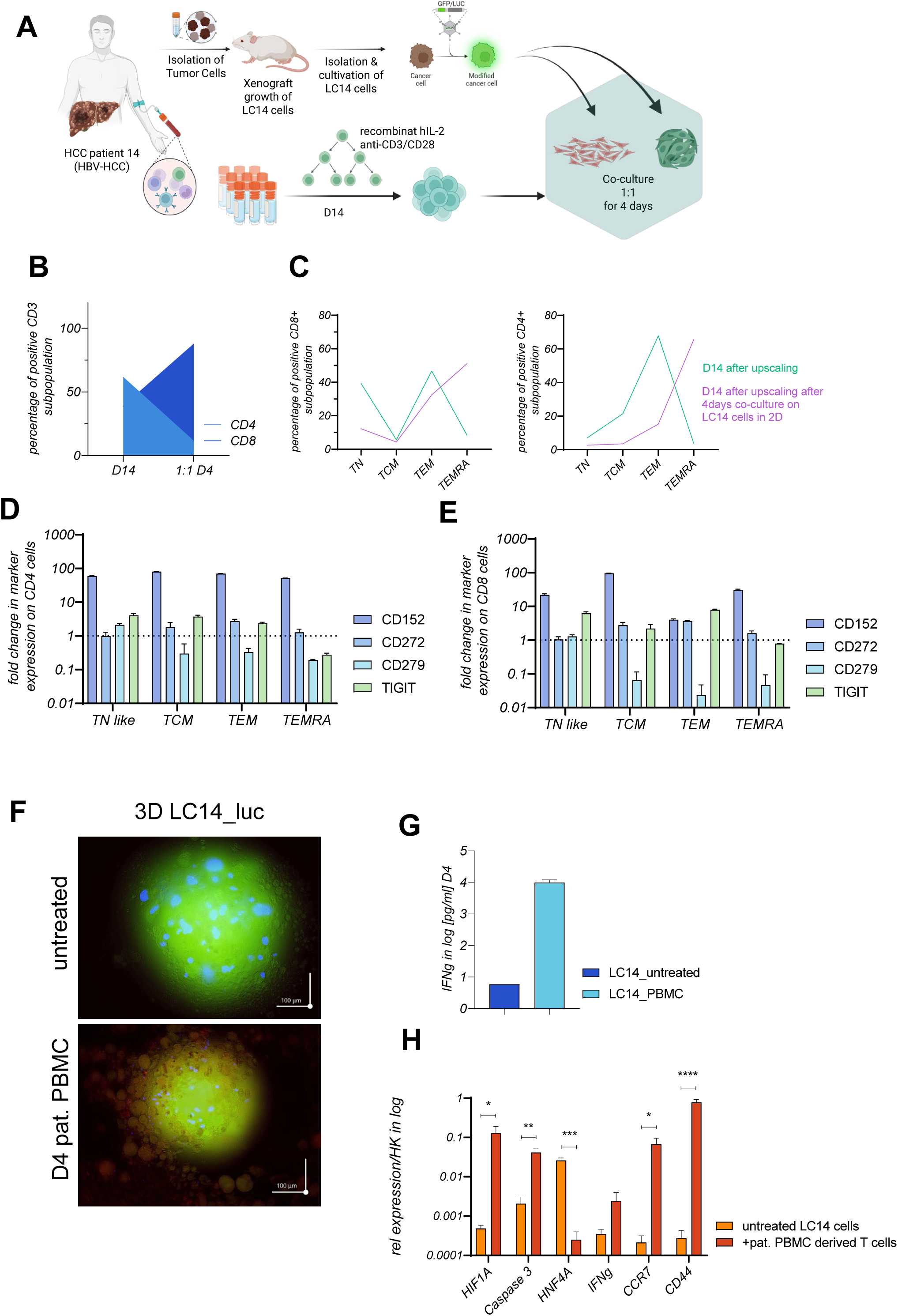
Co-culture of up-scaled HBV patient-derived T-cells with patient-derived HCC cell line. In Figure 5A the autologous experimental setting is visualized. Figure 5B shows the percentual distribution of CD4 and CD8 positive CD3 T-cells from patient#14 before co-culturing and after 4 days of co-culture with the complementary HCC cells from patient#14 (LC14). Figure 5C, displays the T-cell subsets before and after co-culture with LC14 cells (green = initial composition after expansion; purple = composition after 4 days of co-culture). These subsets include TN (naive T-cells), TCM (central memory T-cells), TEM (effector memory T-cells), and TEMRA (effector memory T-cells that re-express CD45RA). Figures 5D and E show the fold change from the indicated inhibitory marker for CD4 (D) and CD8 (E) T-cells, where the initial D0 of co-culture is represented with the dotted line. Figure 5F shows representative captures of LC14 spheroids producing GFP at day 4 of co-culturing with patient#14 derived and expanded PBMCs. In the upper panel, LC14- GFP-spheroid presents the untreated condition and represents the co-culture with patient-derived immune cells in the lower panel. The scale represents 100µm. The patient-derived LC14 cells are visualized in green by stable transfection of GFP-producing LeGo-vector, and patient-derived PBMC, previously treated with a fluorescence dye, are visualized in red. The capturing process was performed automatically by employing an incubation camber and constant exposure time for each channel. The supernatant of the untreated and treated group, shown in Figure 5F, has been used for IFN-γ detection, shown in Figure 5G. Cytokine level is plotted in log [pg/ml], and the LLoD is visualized as the dotted line in the graph. After co-culture, spheroids were used for RNA extraction and gene expression analysis, as shown in Figure 5H. Indicated genes were normalized against RPL0 and GapDH and plotted on a logarithmic scale.

In both subgroups, the TCMs exhibited the highest inhibition markers (data not shown). In contrast, naïve-like, and TEMRA cells, which generated CD45RA, demonstrated the lowest percentages of the specified inhibitory markers (data not shown). As depicted in **Figure 5F**, the 3D co-culture demonstrates a notable reduction in spheroid size, accompanied by discernible ferroptosis structures. The measurement of IFN-γ in the supernatant of the 3D culture, illustrated in **Figure 5G**, emphasises the efficacy of the cytolytic therapeutic compartment by ferroptosis. Additionally, we isolated RNA from the spheroids following 4 days of co-culture and examined the expression of specified genes, as illustrated in **Figure 5H**. Our findings indicate that oxidative stress was elevated in the treated group, whereas HNF4A was significantly diminished, and CCR7 and CD44 were upregulated, suggesting that the enriched patient-derived PBMCs moved into the spheroid and the LC14 cells shifted from proliferation to evasion. The enhanced expression of Caspase 3 in the spheroid structure signifies the initiation of apoptosis. Consequently, we determined that T-cells from patient #14 were capable of causing cytolysis through IFN-γ, GZMB, and the death receptor ligand, thereby triggering ferroptosis, proptosis and apoptosis.

## Discussion

Though effective in hematologic cancers, T-cell-based therapies face significant barriers in HCC due to the highly immunosuppressive tumor microenvironment (TME), restricted T-cell infiltration, and a tendency toward T-cell exhaustion. Particularly in HBV-related hepatocellular carcinoma, persistent inflammation and immunological checkpoint expression promote individual immune evasion strategies and hinder T-cell function [8, 18, 19]. We first tackled the significant challenge in solid tumor immunotherapy: achieving cytotoxic efficacy while avoiding excessive development into senescent TEMRA cells, which may restrict long-term functioning. Secondly, by using HBV-related HCC patient-derived cancer cells from the same individuum, we could mimic the individual suppressive TME, allowing us to examine the functionality and durability of enriched T-cells following expansion.

We effectively enhance effector memory T-cells (TEM) without excessive differentiation into senescent TEMRA cells. The preservation of TEM cells endures anticancer efficacy in solid tumors [6]. TEM cells are crucial for fast antigen response and long-term memory retention, improving persistence in difficult tumor microenvironments [6]. Our findings enhance this comprehension by validating that TEM subsets are efficiently grown in an immunosuppressive milieu, offering additional proof that TEM cells are essential for enduring cytotoxic responses in viral-induced HCC. In line with others, which indicated that TEMRA cells are pivotal in acute cytotoxic reactions, we found a rapid differentiation into TEMRA cells when presenting liver cancer cells from the same patient, in-vitro [5]. Moreover, the observed cell killing points out their distinct cytotoxic activity and role in rapid tumor cell eradication, potentially benefiting clinical applications necessitating short-term, robust cytotoxicity.

On the other hand, we found partially and heterogeneously elevated expressions of inhibitory markers on the expanded T-cells from HCC patients. Recently, checkpoint inhibition has been proposed to mitigate T-cell depletion in solid tumors, extending their functionality within the tumor microenvironment (TME) [1]. Combining therapeutic T-cell compartments with checkpoint inhibitors is essential for sustaining long-term efficacy in HCC [20]. Following this suggestion, elucidating the individual inhibitory signatures and integrating counteracting checkpoint inhibitors into the treatment regime augments the cytotoxic capabilities of the therapeutic T-cell compartment in treating HCC.

## Author contributions

JK initiated and supervised the research study; JK, LS and MD designed the experiments; AH and KS supervised the collection of human samples; JK, LS, GR, MG and TV conducted experiments and acquired data; JK, GR, GM analysed data; JK, LS, SL, SP, MD wrote the manuscript. All authors discussed the data and corrected the manuscript. All authors had access to the study data and had reviewed and approved the final manuscript.

## Acknowledgements

We thank Tobias Gosau and Ursula Mueller for their excellent work. We thank Claudia Dettmer, Eileen Maly, and Karina Börner for their excellent technical assistance. We thank Kristoffer Riecken for providing the LeGO-vector, which was used to transduce LC14 cells stably.

## Aberration

HCC: Hepatocellular carcinoma
HBV: Hepatitis B virus
TME: Tumor microenvironment
CD152: Cytotoxic T-lymphocyte-associated protein 4
CD272: B- and T-lymphocyte attenuator
CD279: Programmed cell death protein 1
TIGIT: T-cell immunoreceptor with Ig and ITIM domains
LC14: patient-derived cell line 14
TN: naïve like T-cells
TCM: central memory T-cells
TEM: effector memory T-cells
TEMRA: Effector memory T-cells expressing CD45RA

## Literature

1. Albarrán, V., et al., Adoptive T-cell therapy for solid tumors: current landscape and future challenges. Frontiers in Immunology, 2024. 15.

2. Yan, T., L. Zhu, and J. Chen, Current advances and challenges in CAR T-Cell therapy for solid tumors: tumor-associated antigens and the tumor microenvironment. Experimental Hematology & Oncology, 2023. 12(1): p. 14.

3. de Miguel, M., et al., T-cell–engaging Therapy for Solid Tumors. Clinical Cancer Research, 2021. 27(6): p. 1595–1603.

4. Klebanoff, C.A., L. Gattinoni, and N.P. Restifo, CD8+ T-cell memory in tumor immunology and immunotherapy. Immunol Rev, 2006. 211: p. 214–24.

5. Golubovskaya, V. and L. Wu, Different Subsets of T-cells, Memory, Effector Functions, and CAR-T Immunotherapy. Cancers, 2016. 8(3): p. 36.

6. Liu, Q., Z. Sun, and L. Chen, Memory T-cells: strategies for optimising tumor immunotherapy. Protein & Cell, 2020. 11(8): p. 549–564.

7. Martin, M.D. and V.P. Badovinac, Defining Memory CD8 T-cell. Frontiers in Immunology, 2018. 9.

8. Mittra, S., S.M. Harding, and S.M. Kaech, Memory T-cells in the Immunoprevention of Cancer: A Switch from Therapeutic to Prophylactic Approaches. The Journal of Immunology, 2023. 211(6): p. 907–916.

9. Scirgolea, C., et al., NaCl enhances CD8+ T cell effector functions in cancer immunotherapy. Nature Immunology, 2024. 25(10): p. 1845–1857.

10. Yao, Z., et al., Focusing on CD8+ T-cell phenotypes: improving solid tumor therapy. Journal of Experimental & Clinical Cancer Research, 2024. 43(1): p. 266.

11. Ball, K., et al., Strategies for clinical dose optimisation of T-cell-engaging therapies in oncology. MAbs, 2023. 15(1): p. 2181016.

12. Guizhen, Z., et al., The tumor microenvironment of hepatocellular carcinoma and its targeting strategy by CAR-T-cell immunotherapy. Front Endocrinol (Lausanne), 2022. 13: p. 918869.

13. Song, G., et al., Global immune characterisation of HBV/HCV-related hepatocellular carcinoma identifies macrophage and T-cell subsets associated with disease progression. Cell Discov, 2020. 6(1): p. 90.

14. Broholm, M., et al., The Adaptive Immune Response in Hepatitis B Virus-Associated Hepatocellular Carcinoma Is Characterised by Dysfunctional and Exhausted HBV-Specific T-cells. Viruses, 2024. 16(5).

15. Staffeldt, L., et al., Generating Patient-Derived HCC Cell Lines Suitable for Predictive In Vitro and In Vivo Drug Screening by Orthotopic Transplantation. Cells, 2023. 13(1).

16. Ludwig, J. and M. Hirschel, Methods and Process Optimisation for Large-Scale CAR T Expansion Using the G-Rex Cell Culture Platform. Methods Mol Biol, 2020. 2086: p. 165–177.

17. Allweiss, L., et al., Immune cell responses are not required to induce substantial hepatitis B virus antigen decline during pegylated interferon-alpha administration. J Hepatol, 2014. 60(3): p. 500–7.

18. Shen, K.Y., et al., Immunosuppressive tumor microenvironment and immunotherapy of hepatocellular carcinoma: current status and prospectives. J Hematol Oncol, 2024. 17(1): p. 25.

19. Raskov, H., et al., Cytotoxic CD8+ T-cells in cancer and cancer immunotherapy. British Journal of Cancer, 2021. 124(2): p. 359–367.

20. Rimassa, L., R.S. Finn, and B. Sangro, Combination immunotherapy for hepatocellular carcinoma. Journal of Hepatology, 2023. 79(2): p. 506–515.

