## Supplementary Table 1 for "Ex-vivo expansion of patient-derived PBMCs preferentially results in effector memory T-cell proliferation with restored autologous efficiency in HBV-HCC"

**Supplementary Tabel 1: FACS Antibody list**

| Panel | Marker | Fluorochrome |
|---|---|---|
| S10 | CCR7 | BB515 |
| | CD152 | PE |
| | CD272 | BV711 |
| | CD279 | PE-Cy7 |
| | CD3 | PerCP-Cy5.5 |
| | CD4 | APC-H7 |
| | CD45 | BUV805 |
| | CD45RA | BV605 |
| | CD8a | AF700 |
| | TIGIT | V450 |
| S9 | CD3 | BV395 |
| | CD45 | BV480 |
| | HLA-DR | BV605 |
| | CD16 | BV650 |
| | CD4 | BV786 |
| | CD56 | PE |
| | CD8 | PerCP-Cy5.5 |
| | CD19 | BUV737 |
