## Supplementary Table 2 for "Ex-vivo expansion of patient-derived PBMCs preferentially results in effector memory T-cell proliferation with restored autologous efficiency in HBV-HCC"

**Supplementary Tabel 2: TaqMan assay list**

| gene name | Assay-ID |
|---|---|
| GapDH | Hs99999905_M1 |
| RPL0 | Hs00420895_gH |
| IFNgamma | Hs00989291_m1 |
| IL2 | Hs00174114_m1 |
| MKi67 | Hs04260396_g1 |
| Caspase 3 | Hs00234387_m1 |
| IL-6 | Hs00985639_m1 |
| IL-10 | Hs99999035_m1 |
| GZMB | Hs01554355_m1 |
| HBV S | Pa03453405_s1 |
| CXCR1 | Hs04187059_m1 |
| CCR9 | Hs01890924_s1 |
| CXCR4 | Hs00607978_s1 |
| CCR7 | HS01013469_m1 |
| ILA17 | HS00174383_m1 |
| CXCR3 | HS01847760_s1 |
| CCR5 | Hs99999149_s1 |
| CCR3 | Hs00266213_s1 |
| CCR2 | Hs00356601_m1 |
| HNF4A | Hs00230853_m1 |
| HIF1A | Hs00936375_m1 |
| CD44 | Hs01075864_m1 |
| CD274 | Hs00204257_m1 |
