## Supplementary Figure 1 for "Ex-vivo expansion of patient-derived PBMCs preferentially results in effector memory T-cell proliferation with restored autologous efficiency in HBV-HCC"

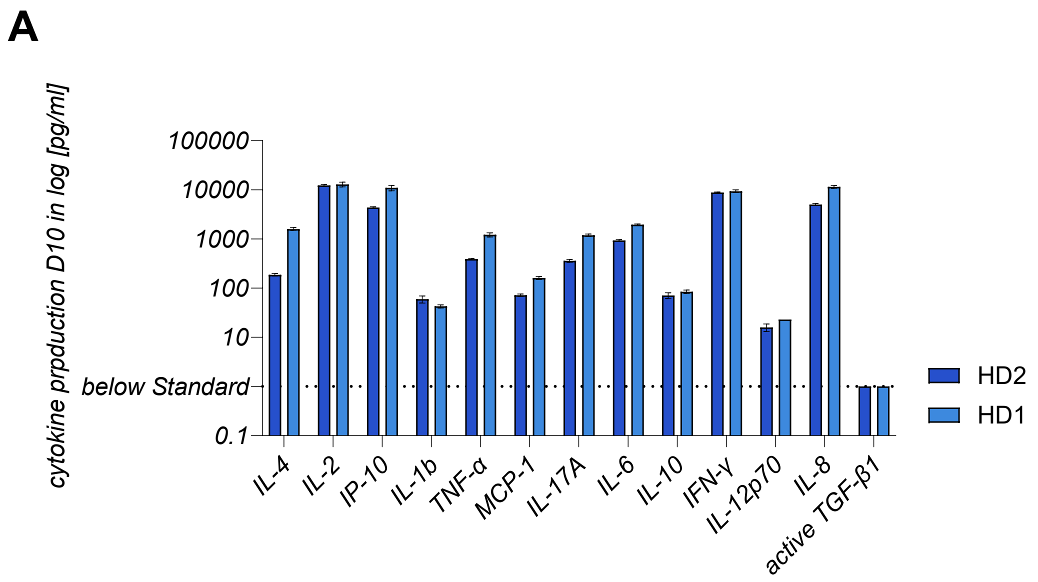

### Supplementary Figure Legends

#### Supplementary Figure 1: Cytokine and chemokine level of HD-derived T cells

In the panel A, the cytokine and chemokine expression measured in the supernatant was acquired on the day 10 of expansion with CD28/CD3-bead based selection and IL-2 application. Supernatant was used for n = 2 technical replicates to be analyzed employing the Legendplex human inflammatory kit, followed by the according quantification software, as described in the method section.
